# Genetic design automation for autonomous formation of multicellular shapes from a single cell progenitor

**DOI:** 10.1101/807107

**Authors:** Evan Appleton, Noushin Mehdipour, Tristan Daifuku, Demarcus Briers, Iman Haghighi, Michael Moret, George Chao, Timothy Wannier, Anush Chiappino-Pepe, Jeremy Huang, Calin Belta, George Church

## Abstract

Multi-cellular organisms originate from a single cell, ultimately giving rise to mature organisms of heterogeneous cell type composition in complex structures. Recent work in the areas of stem cell biology and tissue engineering have laid major groundwork in the ability to convert certain types of cells into other types, but there has been limited progress in the ability to control the morphology of cellular masses as they grow. Contemporary approaches to this problem have included the use of artificial scaffolds, 3D bioprinting, and complex media formulations, however, there are no existing approaches to controlling this process purely through genetics and from a single-cell starting point. Here we describe a computer-aided design approach for designing recombinase-based genetic circuits for controlling the formation of multi-cellular masses into arbitrary shapes in human cells.

## 1 Introduction

Developmental biology is the study of the process by which multi-cellular organisms grow and differentiate. All of these processes can be traced back to a single ‘totipotent stem cell’ from which all differentiated cells originate. These single cells robustly divide and differentiate with impeccable accuracy based on the genetics of this original cell and its surrounding environment. While it is understood that this process happens in different ways for each organism, many of the critical steps along the way are not well understood. Furthermore, the ability to coerce cells to develop in ways outside of the normal developmental context (i.e. stem cell engineering) is relatively nascent. The fields of stem cell biology, regenerative biology, and tissue engineering have long-term goals for using the knowledge of how cells and organisms develop to create novel solutions for disease modeling, drug testing, organ replacement, among other applications [1].

While most of these applications are still currently science-fiction, there have been significant advancements in the basic foundations on which these goals could be accomplished. Namely, the field of stem cell biology has been very active in producing methods for differentiating stem cells into other types of cells following the breakthrough creation of induced pluripotent stem cells (iPSCs) [2]. These methods span from using over-expression of transcription factors [3], to recapitulating the natural developmental environment with engineered surface conditions and complex media with growth factors [4], to using 3D bioprinters to place different types of cells in specific locations [5]. Recently, these types of approaches have even been used to engineer small organ-resembling masses called ‘organoids’ [1, 6].

To date, there have been clear advances in the ability to differentiate cells into different types, but there is still a major struggle to steer cells into forming structured cellular masses that perform a systematic organ-like function. Furthermore, repeatability of the formation of structures as cells grow has been problematic. Recent efforts in human stem cell biology have demonstrated the successful recapitulation of development of human embryos *in vitro* up to 13 days post-fertilization [7, 8] but again, there is no control over the morphology formed at this small scale. 3D bioprinting is one proposed solution to this problem, but this does not always integrate well with genetic processes because placing different types of cells physically next to each other does not necessarily mean that the cells will behave as intended, had they been produced via cell divisions and genetic control.

One possible solution to these obstacles would be to create genetics-controlled methods to control both cellular identity and morphology. In such a solution, structures could form seamlessly as cells divide, with identity changes also controlled genetically. This approach would require synthetic DNA to force cells to form structures and change identity. Fortunately, there are many examples of using genetics to force cell type conversions - the primary knowledge gap is how to use genetics to inform shape formation. Some preliminary work has been done in this area using cadherin surface binding proteins and cell-signaling [9], but this problem is still relatively unexplored.

In order to build genetic solutions to both cell identity control and shape control, one would need to integrate recent advances in the areas of synthetic biology, modeling, and computer-aided design (CAD) to build ‘genetic circuits’ for this purpose. In synthetic biology, genetic circuits are compositions of DNA fragments that each independently produce a cellular function (i.e. transcription, translation, etc.). When stitched together, these circuits can perform a more sophisticated function [10, 11, 12, 13]. Over the past 20 years, many genetic circuit design components have been thoroughly characterized and design principles have been established [14]. More recently, design automation and modeling have been used to automatically design genetic circuits from a library of characterized components [15, 16], giving rise to a variety of successful large, complex constructs in a variety of organisms [17, 18, 19, 20].

Here we describe a computer-aided design software tool, called *CellArchitect*, that uses design automation approaches to design genetic circuits to control the shape formation of multi-cellular masses originating from a single cell (**Figure 1**). To do this, we first breakdown this large problem into smaller sub-problems related to cellular development. Specifically, we identified and solved the following sub-problems: selection of orthogonal cell surface binders to form a shape, definition of a ‘developmental tree’ that maps a cellular population back to its original progenitor, automated design of recombinase-based genetic circuits to control when during development specific genes are expressed, and modeling of cell growth with these genetic circuits integrated into their genome. We make the argument that even if our current solutions are not optimal for forming all shapes, that a computer-aided design solution is needed to attempt these types of problems — human cells divide and differentiate so slowly that it would take an intractable amount of time and money to attempt to solve these problems through trial-and-error. As a proof of principle, we focus on a few small examples of shapes that could be formed with these tools and the sub-problem solutions for their respective genetic circuit design process. Since our input to these tools is a shape and the output is a DNA sequence, these circuits can be directly synthesized and tested in the laboratory.

**Figure 1:**
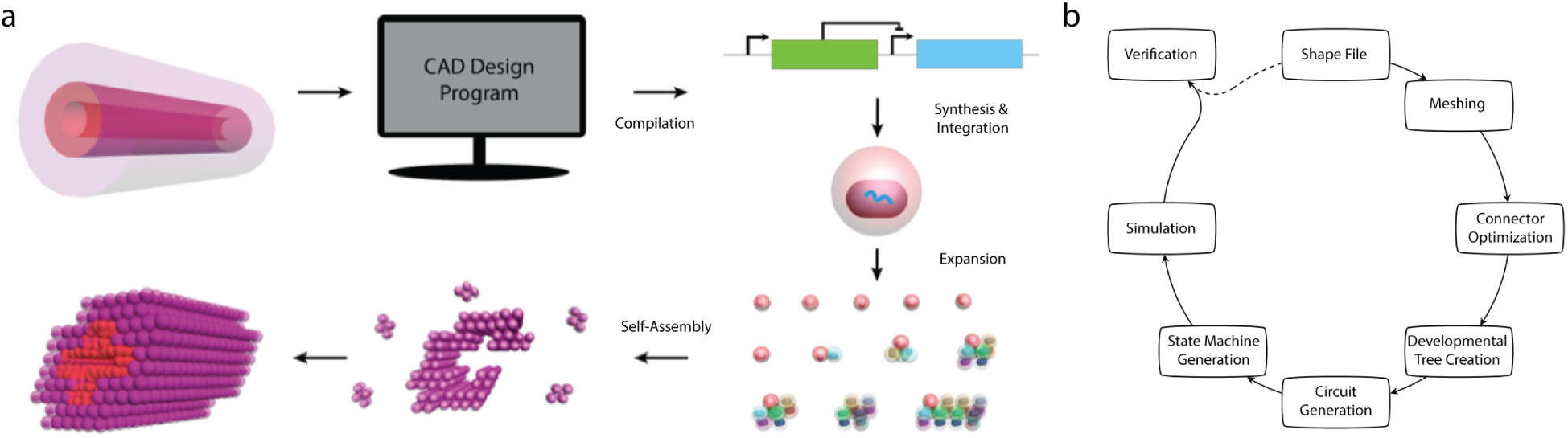
Goal and overview of the software tool process. a. The CAD software inputs a shape file, compiles it to a genetic circuit, which is then synthesized and integrated into the genome of a cell. This cell expands and self-assembles into the desired shape. b. The CAD software sub-problems and how they link together.

## 2 Results

### 2.1 Decomposing large shapes into cellular blocks

The first sub-problem to address when solving how to form large structures of many cells, is how to break these cells into more easily built sub-structures (**Figure 1**). In our software framework, we choose to solve this problem using an approach called meshing - subdividing continuous geometric space into discrete geometric and topological units. This strategy is used in engineering fields for different types of analyses [21, 22], but in our case we use it to divide tessellate 3D-space into discrete tetrahedrons (‘tets’) and cubes (‘quads’). We result in shapes made of these geometric elements that resemble the original shape in an arbitrary amount of granularity (**Figure 2**). We choose to represent larger objects as compositions of tets and quads because we have identified these as achievable building blocks from the 4-cell and 8-cell stage of early embryonic development. The overall strategy is to represent large shapes as sets of ‘cellular blocks’ that can be individually created and stuck together with a calculated choice of orthogonal surface-binding proteins [23, 24].

**Figure 2:**
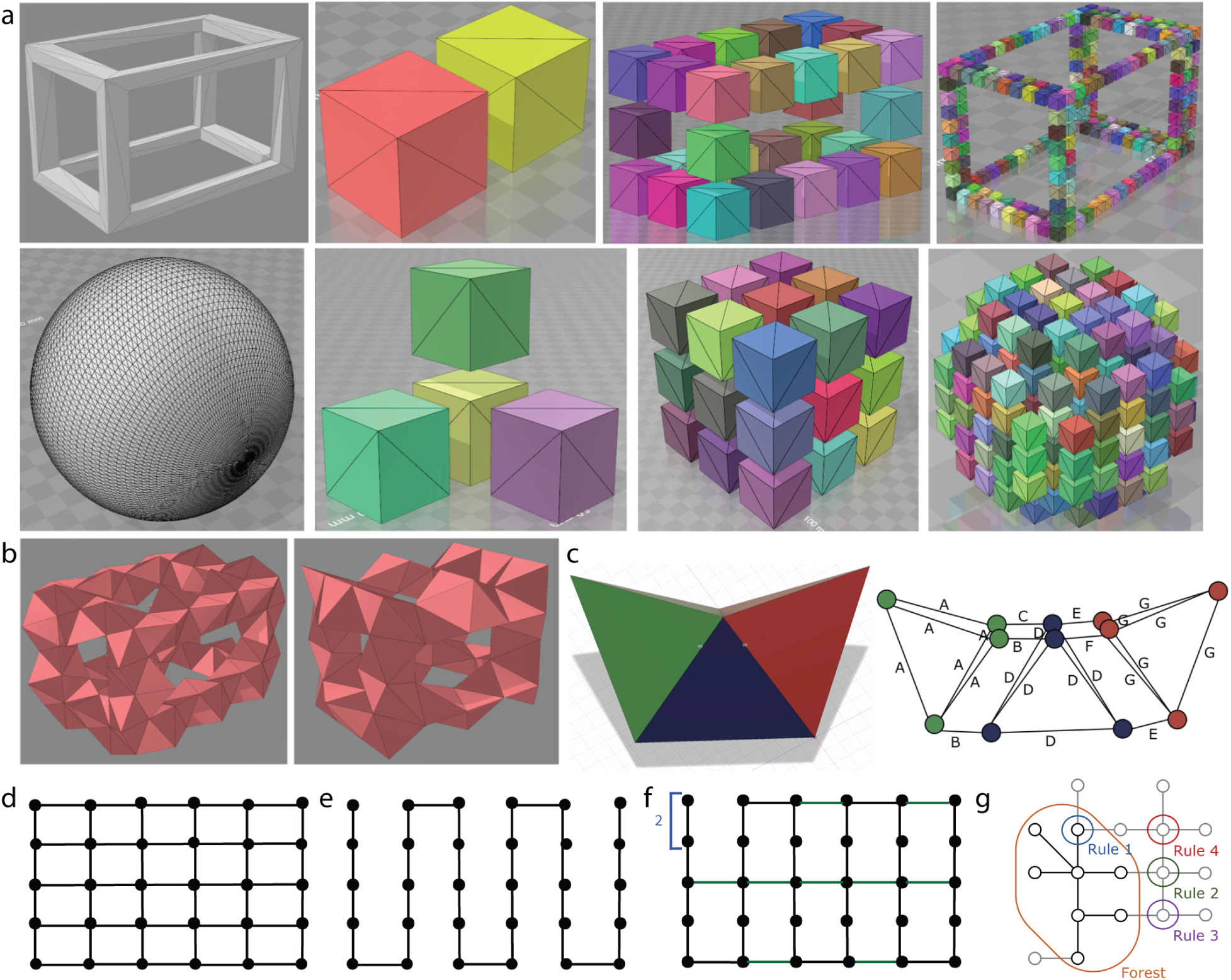
Meshing and connection optimization. a. Two example shapes rendered from an STL file and how they are meshed into cubic approximations of variable numbers of cells. The top row shows how a hollow box would be meshed into a 16-, 256-, and 2048-cell approximation as quads. The bottom row shows how a sphere would be meshed into a 32-, 256-, and 2048-cell approximation as quads. b. The hollow box from ‘a.’ meshed into tets using the ‘Irish Bubble’ meshing algorithm. c. A toy shape can be meshed into 3 tets that interface one another composed of 12 cells. Using the MLST algorithm, we can determine where linkages need to be placed to hold the shape together. d. A fully connected graph where all possible linkages between components are preserved. e. The MST algorithm starts with a fully connected graph and finds a random path of linkages to hold all of the nodes together. f. To reduce floppiness of an MST, we traverse the MST graph adding connections to nodes with only 2 connections. g. The MLST algorithm uses prioritized heuristics to traverse the graph and add linkages based on these heuristics.

The input file to this part of the workflow (and the workflow more generally) is an STL file [25], a standardized CAD file format in either ASCII or binary. This ‘shape file’ is the starting point from which the genetic circuit is determined, and is used a comparison to the multi-cellular results produced by the simulator and experimentally acquired microscopy data. These comparisons give a statistical metric of success for candidate genetic circuits and also a quantitative measure of success of experiments compared to what is desired in the shape specification step.

#### 2.1.1 Creating cellular meshes

In order to break up user-provided target shapes into blocks of cells, we initially turned to the world of shape meshing algorithms. However, canonical shape meshing algorithms proved ill-suited to our application, as they generally prioritize shape fidelity over block consistency – that is, in order to achieve a mesh that strictly matches the original shape, the algorithms will choose blocks that are significantly deformed, or are of disparate volumes. Because we chose to model K562 cells as spheres of identical volumes, it was important to find a meshing technique that would instead prioritize generating blocks of regular shape and size.

For hexahedral meshing, this was as simple as overlaying a 3-dimensional grid of cubes over the target shape and retaining only those cubes that fell within the shape, thus ensuring that every cube was identical (**Figure 2a**). A similar method was used to create a tetrahedral mesh but, as regular tetrahedra do not tessellate space, it was necessary to instead overlay a tessellation of near-regular tetrahedra. We therefore implemented the “Irish” Tessellation [26] which tessellates space using 3 types of tetrahedra, the largest of which has a volume less than 13% larger than that of the smallest, and all of which have dihedral angles between 53.1 and 78.5 (a regular tetrahedron has a dihedral angle of 70.5) (**Figure 2b**). As with the hexahedral meshing, only tetrahedra that fall within the shape are retained.

#### 2.1.2 Minimization of orthogonal surface binding proteins

Once a cellular mesh has been created, we aim to minimize the number of orthogonal surface-binding proteins required to hold the desired shape. This is done because the number of well characterized orthogonal surface protein binders currently known is relatively small. This step takes a mesh for the desired shape as input and outputs a list of cell-cell connections required for the shape to be formed. We solve this problem with two alternative type of applied algorithms: a Maximum Leaf Spanning Tree (MLST) algorithm [27] and a Minimum Spanning Tree (MST) algorithm [28].

The first algorithm we use to minimize connections in a mesh is MLST [27]. The algorithm relies on maintaining lists of nodes (i.e. cell in a cellular block and its neighboring cells in the mesh) that qualify for different rules in a forest (i.e. MLST we are building):

Rule 1 Any node in the forest with at least two neighbors not in the forest.
Rule 2 Any node not in the forest that has at least one neighbor in the forest and at least three neighbors not in the forest.
Rule 3 Any node not in the forest with at least one neighbor in the forest and exactly two neighbors not in the forest.
Rule 4 Any node not in the forest with at least three neighbors also not in the forest.

Starting with a single seed node (i.e. cell in a cellular block and its neighboring cells in the mesh), the algorithm “expands” the forest to add nodes, prioritizing Rule 1 over Rule 2 over Rule 3 over Rule 4 (**Figure 2g**). The algorithm expands the forest until there are no more nodes in the rule lists, then looks for any node in the forest that has neighbors outside the forest and adds those neighbors. Next a breadth first search (BFS) on the forest is performed on the forest to determine if all nodes in the forest are connected. If they are not, the algorithm adds edges from nodes that are connected to the node BFS was run initiated at to the nodes that are not connected until the tree is connected. This algorithm generates a tree that maximizes the number of nodes (blocks) with only one connection, and can serve as a starting point for choosing block connections. It is hypothesized that having more leaves may lead to greater shape fidelity, as it would likely lead to a smaller, more dense, core shape. Moreover, when cell-surface binding proteins need to be reused within a shape, leaf blocks are logical candidates because they can all be identical.

The second algorithm we use to minimize cell block connections is a MST [28]. This algorithm finds a generic minimum spanning tree from an input 3D cell-block mesh mapped to a 2D graph. We then take this minimal graph and attempt to reduce “floppiness” by adding edges to minimize the number of nodes with only two connections. The algorithm begins by identifying problem nodes - the nodes with only 2 connections. It then considers all edges it could add to the graph to remove the problem (it identifies these options by comparing the original graph of all possible connections to the MST). Each edge is assigned a score. Edges that solve 2 problem nodes at once receive higher scores. Edges are penalized if they would inhibit other edges that solve 2 problems since we only wish to add one edge per problem node (this stipulation could be revisited). They are also penalized if they connect to leaves. Once these scores are assigned, the list of potential edges is sorted by their scores. Solution edges are added to the graph until there are no more potential edges or all of the problem nodes have been resolved (**Figure 2d-f**).

These two algorithms result in alternative connectivity solutions for the same input shape. Currently, the solution with the fewest required sets of orthogonal surface binding pairs is selected, but the algorithm that produces this answer is subject to the input shape. Finally, this graph is traversed and results in a list of cellular blocks, where each cell is required to have specific binding proteins.

### 2.2 Constructing a developmental tree from a list of cellular blocks

After determining a set of cellular blocks that could form and hold a shape, we must determine how these final cells will be formed from a single progenitor cell. To solve this problem we invented the idea of a ‘developmental tree’ - a map back from which all final cells arise from the original cell. This map tells us which progenitor cells must undergo certain biological processes en route to a final set of cells. The key behaviors that we consider in the context of this problem are asymmetrical cell divisions, cell surface binding protein expression, cell-cycle arrest and apoptosis. Asymmetrical cell division is key because unique shapes require heterogeneous cell populations, cell-surface binding proteins to form the shape, cell cycle arrest to stop cells dividing to hold a structure, and apoptosis to remove extra cells that are not needed for a structure. To solve this problem, we created an algorithm that inputs a set of cell-cell connections and outputs a tree illustrating which cells must exist over time to give rise to the correct final set of cells to create the input graph. It determines which surface proteins need to be expressed by each cell, then groups cells based on these proteins and determines a pathway to go from one cell to the final population.

The module begins by going through each block in the graph and determining the surface binders that need to be expressed by each of its cells. Tet blocks are bound together by a single homodimer. Quad blocks are bound together with three heterodimers: one binder to make a trigonal planar shape with four cells, another to make a second trigonal planar shape with the other four cells, and then the third binder to interlock the 3 outside cells of each trigonal planar shape (**Figure 3**). Connections between hex blocks are made using 2 homodimers, one at each end of a diagonal across the face of the cube. Care is taken to ensure that the orientations of the blocks relative to each other is preserved: this is achieved via a lookup table that correctly matches the corners of the two blocks. For tet blocks, one homodimer and one heterodimer are used to bind blocks, the heterodimer at two corners and the homodimer at the third. The relative orientation between two blocks is again preserved: binders are assigned to specific corners depending on which face is in play. In both the tet and hex cases, multiple binders are used to try to prevent rotation of blocks against each other.

**Figure 3:**
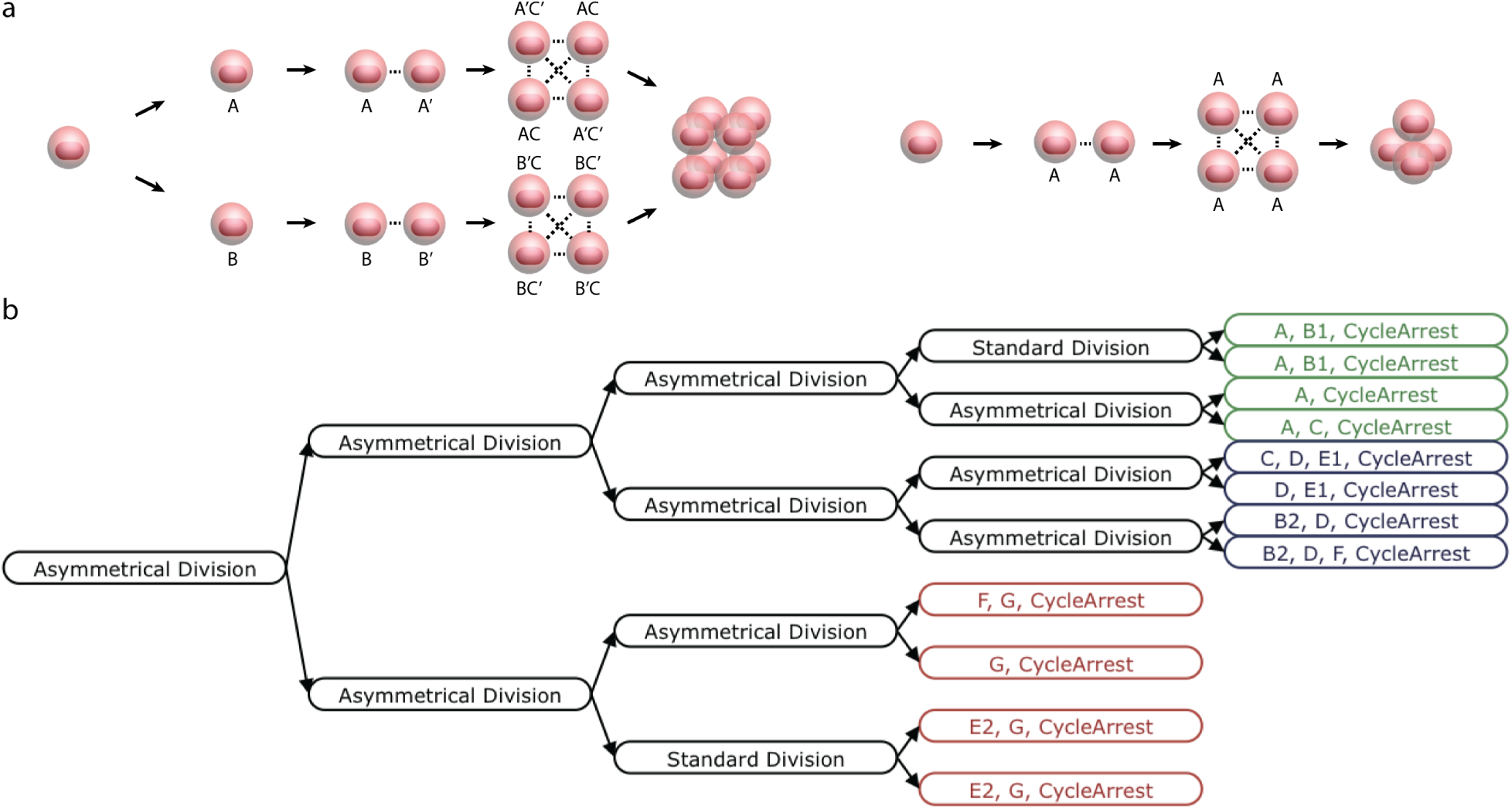
Developmental trees are produced from lists of blocks and connections they must form. a. Each tet block is made from a single homodimer (protein A) and each quad block from three orthogonal heterodimer pairs (proteins A/A’, B/B’, and C/C’). b. Starting from the final cells that must exist in a shape, a binary tree can be created to determine how many divisions must happen to form all of the blocks and optimize where asymmetrical divisions must occur.

Once all the binders have been assigned, the cells are sorted by the proteins they are expressing. Cells are added to the tree (stored in a list) starting at the bottom layer. If the final number of required cells is not a power of two, there will be two bottom layers. Note that any number of cells *n* ∈ ℕ can be obtained by building a perfect binary tree of height *K* ∈ ℕ where 2^*k−*1^ ≤ *n <* 2^*k*^, then having *n* − 2^*k−*1^ cells in the bottom layer of the tree divide once more, so we need at most two bottom layers. Cells at the bottom layer are then merged in pairs to create the preceding layer. The merging process determines if the two cells are identical, and can therefore be generated by a standard division, or if they their proteins differ, and must therefore be generated by an asymmetrical division. This process is repeated until the top of the tree is reached. If there are two bottom layers, then the second lowest level will contain a mix of cells that must divide again and cells that are in their final state. As each layer is created, it is appended in order, cell by cell, to the list containing the developmental tree, with each cell containing a code — −1 for no division, −2 for standard division, and −3 for asymmetrical division — as well as the set of expressed proteins (surface binders and fluorescent proteins) determined previously. The tree in this format can easily be read from right to left to pull out the information on tree structure: the top layer is in the last spot, the second layer is in the second and third to last spots, and so on, with each layer being read internally from left to right.

Finally, we perform one final optimization on this partially-complete developmental tree in attempt to maximize similarity of ancestor cells in order to minimize asymmetrical division events. We take a list of sets containing protein names where each set represents a cell and its expressed proteins, and we group cells based on the similarity of their sets of proteins. We begin by calculating the ratio of shared proteins to total proteins between every pair of cells and create an undirected graph where cells are nodes and the ratios are edge weights and then generate a hamiltonian path through the graph, using a greedy approach — we the edge with the highest weight that does not violate the hamiltonian requirements until the path is complete.

### 2.3 Designing a genetic circuit from a developmental tree

To execute the genetic program of a given developmental tree, the cells must have a way in which to program specific genes to become expressed at discrete cell divisions. We choose to control this process through the use of genetic counting circuits. Specifically we use DNA recombinase components to invert and excise DNA [20, 19], cell cycle dependent promoters [29] and degradation tags [30] to have recombination events only once per cell cycle, and cell cycle-arrest [31] and apoptosis-inducing genes [32] (**Figure 4a**). We report our designs for genetic counter constructs that express specific recombinases in each cell cycle and genetic register constructs that manipulate which genes are expressed as a function of which recombinase is present in the cell (**Figure 5**). We describe three general options for counter and register constructs.

**Figure 4:**
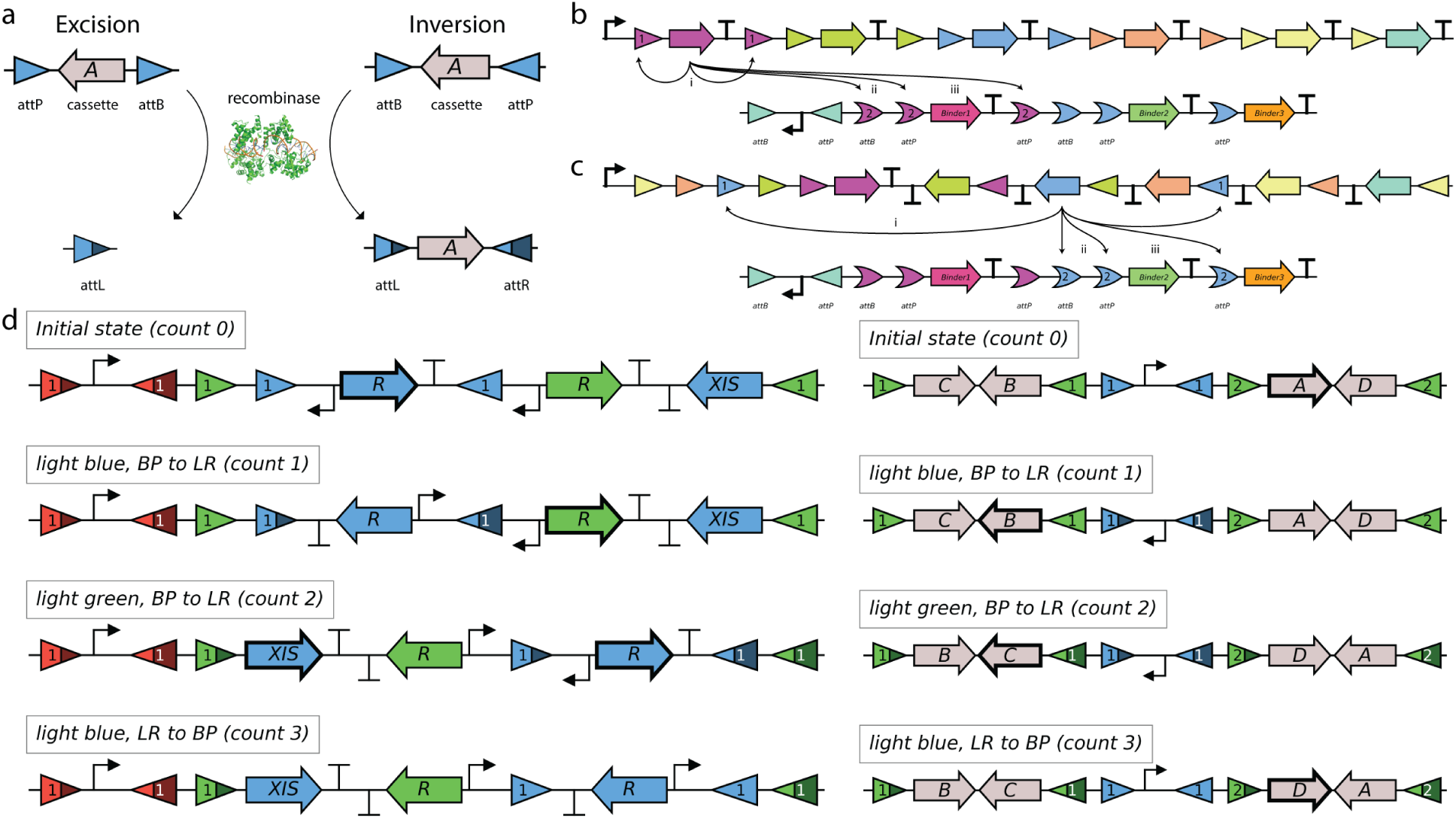
Genetic recombination for genetic circuits that count cell divisions. In this figure, all promoters are cell-cycle dependent promoters that initiate transcription for less time than it takes to produce a mature protein from the transcript it initiates. a. Serine integrases recognize pairs of recognition sites called attB/attP and attL/attR sites. attB/attP sites are converted to attL/attR upon recombination and upon use of an RDF they can be reverted to attB/P. When the recombinase sites are on the same strand the internal DNA is excised and discarded, when the recombinase sites are on opposing strands, the recombinase inverts the DNA in the middle. b. A linear excision counter and register. In the first count, a recombinase expression transcript is initiated and once it is fully mature, it excises regions from both the counter and register (pictured in maroon). In this example, we also show how using orthogonal recombination site pairs (‘1’ and ‘2’) allow us to perform asymmetrical divisions — when the recombinase is expressed, it will perform excision ‘i’, but either ‘ii’ or ‘iii’, not both, since ‘ii’ makese ‘iii’ not possible and vice-versa. After recombination, the next recombinase is exposed for transcription. c. A linear inversion counter and linear excision register. These constructs are similar to linear excision devices, except DNA strands are inverted to advance counts. Asymmetrical division is illustrated with the blue recombinase as per in ‘c’, except that the recombinase will flip the counter DNA and excise the register DNA. d. A binary inversion counter and register. These constructs use inversion like the linear inversion circuits, but also use RDFs (a.k.a. XIS genes) to convert attL/attR sites back to attB/attP sites, again inverting the DNA in the process. Because it uses these RDFs to ‘flip back’ sections of DNA, it counts in binary and can accomplish 2^*n*^ operations per recombinase.

**Figure 5:**
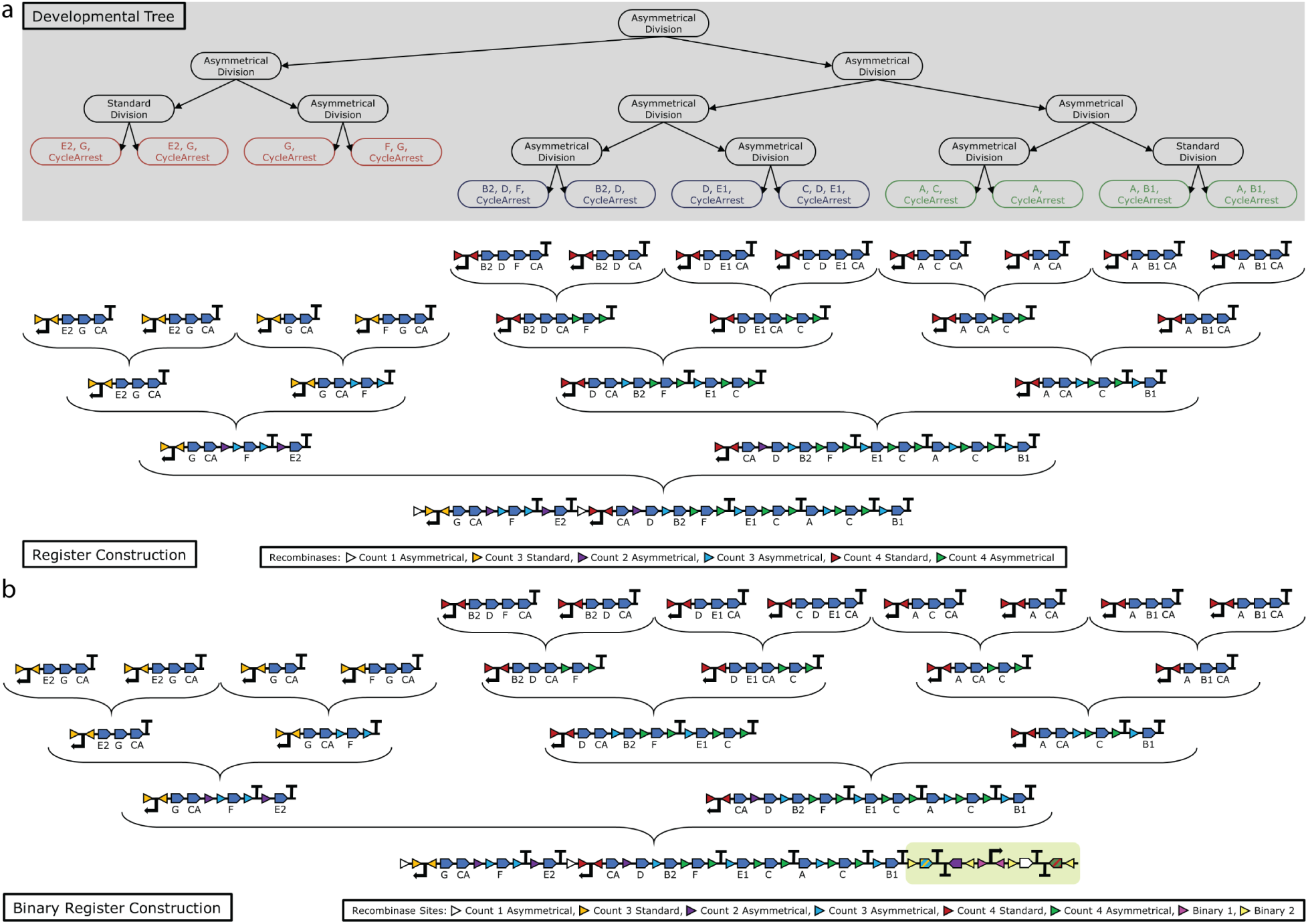
A genetic register is built recursively based on the developmental tree input. This register is built for the 3-tet shape using its developmental tree as input. After optimization to minimize asymmetrical divisions, final expression circuits are given to the final steps and built upwards using asymmetrical division conventions using multi-recombination. a. A linear excision register. built recursively from the ‘leaves’ b. A binary inversion register built recursively from the ‘leaves’. The addition to this register that demonstrates the difference between this and the linear register is highlighted in yellow

#### 2.3.1 A genetic counter circuit that counts cell divisions

The counter construct drives the behavior of the register and can be designed in three ways: linear excision, linear inversion [13], and binary inversion. The linear options are simpler but you can only count as high as the number of orthogonal recombinases you have and the construct’s DNA base length scales linearly with count number (**Figure 5**). To solve this problem, we invented a circuit architecture to count recombination events in a binary manner. The binary counter can count to 2^*n*^ with *n* recombinases. We reason that since a human body has approximately 37 trillion cells, approximately 45 to 50 counts should be an adequate ceiling for counts required for a large hypothetical human structure and therefore linear constructs would need 50 orthogonal recombinases to count that high, and binary constructs only 6.

For the linear inversion architecture, this algorithm puts the first recombinase in the forward orientation following the promoter, and puts all subsequent ones in order in the reverse orientation, with facing recombinase sites radiating outwards (so every count flips a larger piece of DNA, and flips the next recombinase into position after the promoter) (**Figure 4c**). For the linear excision architecture, we have one promoter at the starting position and oriented forward into a gene and terminator that will get excised upon production of the recombinase before the following count (**Figure 5b**).

For the binary counting architecture, we uses a pattern in which n recombinase/reverse-directionality-factor (a.k.a. RDF or XIS) pairs [33] can be used to count 2^*n*^ genetic recombination events. If there is only one recombinase, then we put the first recombinase after the cell cycle dependent promoter. Otherwise, we begin with a forward cycle dependent promoter. Then for each recombinase in the recombinase sequence, we insert a forward attB site immediately after the initial promoter. For the first two recombinases, we then add a reverse cell cycle dependent promoter. For all recombinases, we then put the recombinase gene in the forward direction, then add a forward terminator, then, for all but the first recombinase, we add a reverse terminator followed by the RDF of the previous recombinase in the reverse direction. Finally, we add a reverse attP site for each recombinase (**Figure 4d**).

#### 2.3.2 A genetic register circuit expresses genes of interest at discrete genetic counts

The register constructs have a constitutive promoter and orthogonal pairs of recombination sites to manipulate which genes are transcribed given which recombinase is currently expressed by the counter [34, 19] (**Figure 4d**). An algorithm designs a register circuit based on a developmental tree. This method can create a register compatible with a binary counter architecture or a linear architecture. The code begins by reserving some recombinases for use in the counter, as additional recombinases will generally be required for asymmetrical divisions in the registers. The register architecture is divided into two kinds of sub-registers -‘the final count register’ and the ‘frame register’. The final count register is the same for the two architectures. Note that unlike the rest of the register, this final expression register is expressed on a constitutive promoter as we need the surface proteins to be expressed despite the fact that we have halted the cell cycle.

For the linear registers we create a replica of the counter architecture without the recombinases and replace the recombination sites with alternative, orthogonal recombination sites to complete the frame registers. For the binary architecture, the algorithm takes the log_2_ of the number of counts and rounds up to get the number of recombinases. The frame register for the final count is then determined from the binary tree - an alternating-strand architecture is employed: a central promoter that flips directions every other count with output slots fanning out on either side, counts 1, 4 (rev. strand), 5, 8 (rev. strand), … on the right and counts 2 (rev. strand), 3, 6 (rev. strand), 7, … on the left.

#### 2.3.3 Recursive design of register constructs based on developmental tree

A recursive function then builds the final output register that corresponds to a developmental tree (**Figure 5a**). For the linear registers, the algorithm descends the tree until it reaches the leaves, putting in required sites for asymmetrical division as necessary. Once it reaches a leaf, it creates the expression circuit for that cell, consisting of a reverse promoter that will be flipped at the cell’s final count, and the genes the cell must express. Then, on the way back up, it determines if there is redundancy between two daughter cells (if they are expressing some of the same genes). If so, it moves those genes outside of the asymmetrical division recombinase sites. It also accounts for situations where one daughter cell divides more times than the other (necessitating completely separate circuits, since the final count promoters for those two cell populations must flip on at different counts).

For binary registers, the recursive addition of the final register and asymmetrical components is more complicated — the binary expression register requires the use of nested recombinase site pairs. The algorithm recursively descends through one side of the binary expression register, starting at the largest, exterior pair and calls itself on each interior pair until it reaches one with no interior pair (**Figure 5b**).

This algorithm begins by checking if there is only one count between the current sites. In this case, the count must be the 0th, 4th, 8th, … or 1st, 5th, 9th, … count, as the other counts cannot appear by themselves (they would always be paired with one of the counts listed previously). We can eliminate the tier 1 (the second recombinase, which flips every 4 counts) sites immediately surrounding this count. We must flip this final count if *n* mod 8 ∈ {0 mod 8, 1 mod 8} where *n* is the count in order to ensure that the count is correctly oriented when it should be expressed. Next, we check to see if any of the counts between the current sites actually contain genes. If they do not, then the current sites are not necessary and we can simply put a forward and a reverse terminator in their place (these terminators are necessary to ensure that potential later counts are not expressed prematurely). Otherwise, if we are at the lowest tier (ie we have 2 counts), we use the sites, put the genes for the first count in the forward direction followed by a forward terminator, then add a reverse terminator followed by the genes for the second count in the reverse direction. If we are not at the lowest tier, the function calls itself on the tier below and encloses the output in facing sites.

For our 3-tet shape, the register is formed in 4 recursive stages, starting from the leaves of the last two tetrahedrons to form (**Figure 5a**,**b**). The ‘final registers’ are determined and compiled upwards into the frame registers as dictated by asymmetrical divisions. While the linear and binary register conditions are different, the differences in the final register are relatively minor (**Figure 5b**).

### 2.4 State machines for simulating genetic circuit dynamics

Once a genetic circuit is designed by our software, it should function assuming perfect functionality by its components; however, any synthetic biologist could tell you that this will not be the reality. While the recombinase components are some of the most robust in the synthetic biology world and we can demonstrate very orthogonal binding behavior for some surface binding proteins, these biological components are imperfect. Furthermore, it’s possible that synthetic genetic components will have unexpected interaction with endogenous genetic components. To accommodate for this imperfection, we use the mathematical model of finite state machines as a simulation framework for placing our circuits into living cells (**Figure 5a**). We argue that this framework is fitting for this problem because recombination-based circuits lend perfectly to finite-state behavior — the circuit has one starting condition and can be manipulated via recombinases to another condition where it has other behavior. Based upon experimentally-measured probabilities of a recombination event given the expression of a recombinase, we can assign real probabilities to each of the state transitions. When we couple these recombinaton events to the cell cycle and cell division, we can create a simulated cellular environment that guides individual cell behavior based upon the state of the circuit in each cell and additional environmental factors(**Figure 6a**).

**Figure 6:**
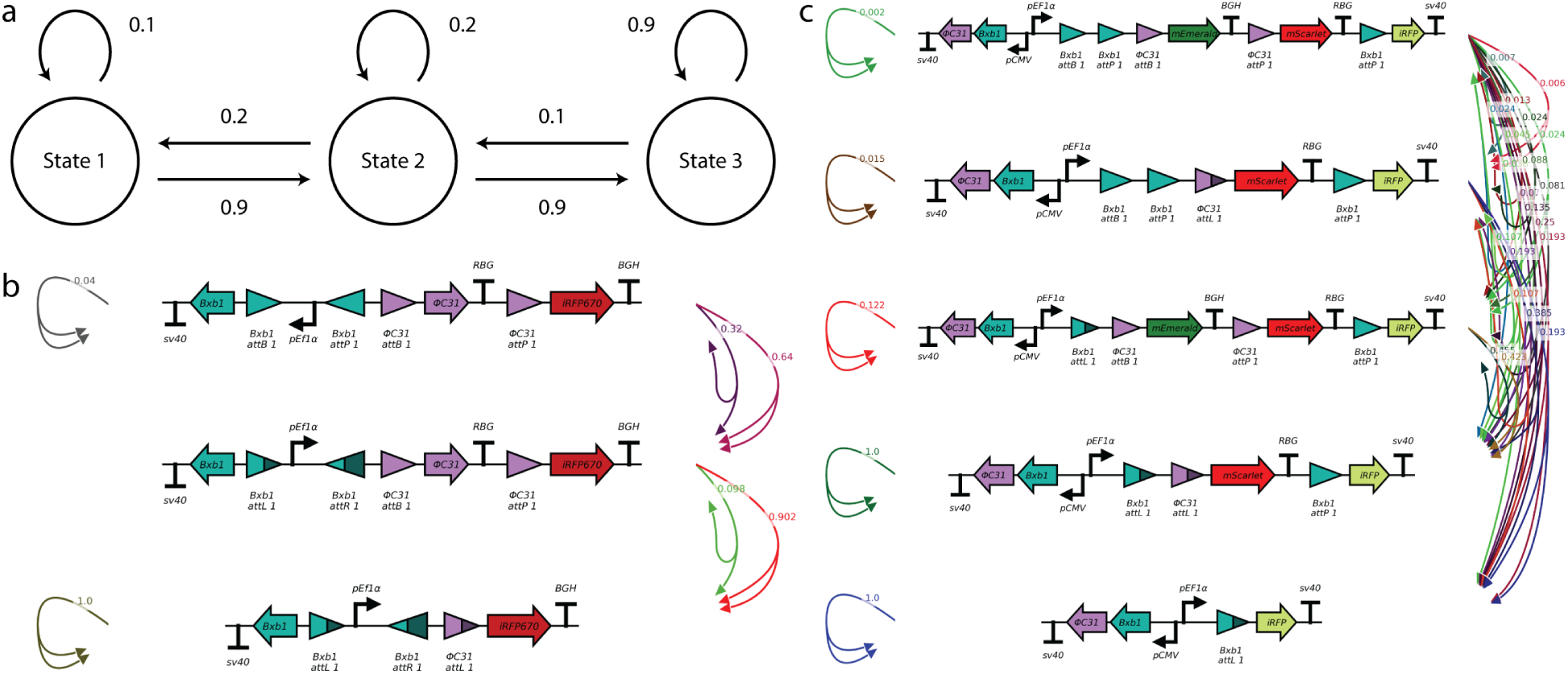
State machines are used in the physical simulation to determine internal cellular conditions. a. An example state machine with 3 states — upon any event, the object has assigned probabilities of transitioning to other states or remaining in the same state. These probabilities all must add up to 1. b. A state machine for a toy recombinase circuit. Since each cell divides into two cell, each of which has a different recombination (state transition) event, state transitions are calculated for pairs of outcomes (i.e. one daughter cell undergoes recombination, the other remains unrecombined at that time). c. A state machine for a counter with five states. We can see that as we consider all of the off-target recombination activities that are possible, the number of edges expands exponentially.

This step inputs a genetic circuit and outputs a state machine reflecting all of the possible recombined states the circuit could end up in (nodes) and the probabilities of each state transition (edges). This state machine is represented slightly differently than the canonical state machine — in our context, we must accommodate for cell division; we have one cell in one particular state that divides into two cells, each with a possible state transition. Therefore, we represent state transitions as pairs of edges as opposed to a single edge. The computing therefore resets for each cell at the beginning of the next cell cycle (**Figure 5b**). As a consequence of this ‘forking’, the state machines for even a relatively simple circuit can balloon in size rather quickly. Furthermore, if we consider rare off-target events, the number of possible states also balloons, and we are left with state machines that increase in complexity exponentially. To reduce complexity of state machines, we eliminate state transitions whose probability falls below a specified threshold and its probability reassigned to other transitions from the original state (methodology explained in greater detail below). This allows us to limit the size of the state machine by decreasing the number of transitions and ignoring states that are very unlikely to ever be reached. The probability for each transition is calculated for one state going to a pair of other states, as we are hoping to tie changes to the circuits to the cell cycle using cycle-dependent promoters and degradation tags and have each circuit operation occur independently in each daughter cell soon after a cell division.

The algorithm to build state machines from a starting circuit starts by scanning the components and identifying the expressed genes and RDFs and checking if the cycle arrest gene is expressed. If the cycle arrest gene is expressed, then no transitions to other states are considered. For states that are not in cycle arrest, we find active recombinase site pairs (i.e. pairs of the same site ID, for the same recombinase, that are of complimentary types (eg attP and attB) and whose recombinase is active). These pairs are stored in a dictionary, and base probabilities for each site pair are looked up (the probability of a recombination event between an attB/P or attL/R site occurring given the expression of a recombinase and/or RDF), then all allowable combinations of site pairs are determined. Each combination will define a state that the current state can transition to. We next calculate the probability of those transitions — these probabilities are directly dependent on which recombinases are active in the cell. It is therefore necessary to calculate the probability of each transition for each possible set *S*_*n*_ of active recombinases such that *S*_*n*_ = *T* ∪ (*n* ∈ *N*) where *T* is the set of all expressed recombinases.

The next step is to threshold transitions that have probabilities that are too low. In order to reduce memory usage, two rounds of thresholding are performed. This is necessary as the second thresholding event considers all size 2 combinations of the transitions that made it through the first thresholding event. The threshold value, however, is meant to be applied to the final size 2 combinations; since we are unable to calculate those immediately, it is necessary to calculate a stand-in probability for each transition for use in the first round of thresholding. The probability is:

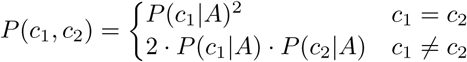

where *c*_1_ and *c*_2_ are combinations of site pairs, *A* is the case where all recombinases are active, which equates to the probability of the first state multiplied by the probability of the second state, times 2 if the states are different since there are then two unique ways to achieve that state pair. This assumes that the efficacy of each recombinase in one daughter cell is independent from the efficacy of that same recombinase in the other daughter cell. The program determine the probabilities of each state-transition option in the following way: the combinations are sorted by the stand-in probabilities, then a binary search is used to determine the first combination whose stand-in probability is above the threshold. Subsequently, alternative combinations are generated for each combination. With large circuits, the combinatorial space of possible state transitions is immense, so when there are greater than 1000 combinations, the program does not intelligently reassign probabilities of problematic combination pairs. Rather it sums the true probabilities of these pairs and reassigns it proportionally to the other pairs according to the following formula: 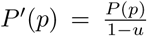 where *p* is a valid pair and *u* is the sum of the cut probability. When there are fewer than 1001 combinations, the program reassigns the probability intelligently using the previously generated lists of alternatives.

The 3-tet shape example has >1000 states for both linear and binary format, so we do not show this example here. Instead we show two example counters (**Figure 6b,c**) to show how the edges can be visualized and that we must accommodate for two cell divisions in this way, as each daughter cell has its own state. All state transitions from one node to all others add up to 1 and when we consider low-likelihood state transitions, the number of possible changes expands very quickly, even for 5 states as opposed to 3.

### 2.5 Modeling multicellular self-assembly in 4D space

Multicellular pattern formation is an emergent behavior in mammals that controls complex behaviors, such as embryonic development, and is tightly regulated by biochemical and mechanical cues in both 3D space and time (we refer to this as a 4D space). Currently, there are several exploratory approaches to induce multi-cellular patterning in mammalian cells using directed self-organization driven by interfacial tension [35], micro-patterned surfaces [36, 37], or biochemical signaling [38]. However, inducing these behaviors requires significant manual intervention from the experimenter or requires the assistance of artificial scaffolds. These approaches that requires manual intervention are difficult to scale up and produce more complex patterning.

In addition to the need for manual intervention, most approaches to control self-organization are driven by trial-and-error design which becomes increasingly difficult as multicelluar systems become more complex. Briers and Libby et. al. [39], demonstrated the ability to combine a 2D Cellular Potts model with Particle Swarm Optimization to automated the design and experimentally validate 2D symmetrical patterns in human induced pluripotent stem cells. To overcome limitations of trail-and-error driven experimentation, and allow more more complex pattern formation, we have developed a computational model to simulate the self-assembly from a single cell into 3D structures with out manual intervention.

Once a state machine has been built for a genetic circuit, we can simulate what would happen to a cell population containing that circuit as it expands into a multi-cellular population. When modeling the role of tissue mechanics (eg cell-cell adhesion, cell migration, cell shape, haptotaxis), the most common agent-based modeling (ABM) frameworks are Cellular Automaton [40], Cellular Potts framework [41], Vertex models, Cell Center-based models (this includes particle models and rigid body models like we use in this paper), and hybrid models that combine aspects of multiple ABM frameworks [42]. We chose to model the programmatic self-assembly of K562 cells as incompressible spheres propelled by Brownian Motion. This modeling framework is appropriate since our experiments involve K562 cells that are cultured in a suspension media lacking extracellular matrix or any stiff surface for self-propulsion by individual cells. For the duration of experiments, the aqueous growth media is autonomously shaken. We model this collision of water molecules with cells as Brownian motion.

In each simulation, a biophysical simulation is run in which the circuit is placed in one starting cell and cells are expanded and genetically modified over time as per the state machine and external conditions. External conditions include the expression of cell surface binding proteins and Brownian motion of cells in a confined space (simulated shaking cell culture). In this case we consider the physical properties of cell line K562 [43] (**Figure 7a**). This cell line is chosen because it is an immortalized blood cell line that naturally has no surface binders. The cell line also has a spherical, generally uniform size — these properties are ideal because they allow us to model the cells as non-adhering uniform spheres for which Brownian motion approximates well when these cells are shaken. In our case Brownian motion is desirable as it increases the frequency of cell-cell contacts needs to self-assemble. Cells that are confined to planar cell migration at the bottom of a dish are extremely unlikely to ever find their correct binding partners in the sparse environment with relatively few cells.

**Figure 7:**
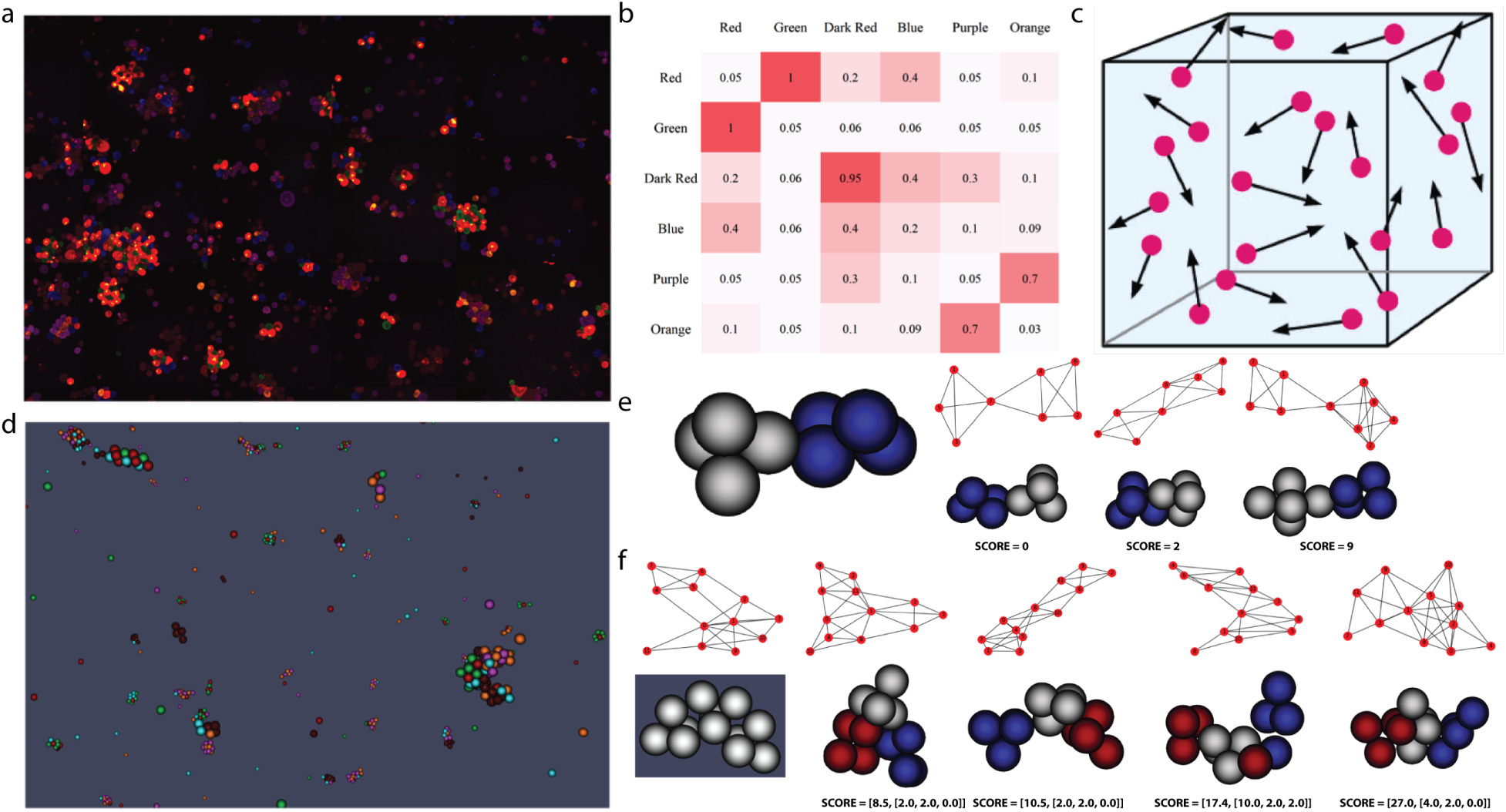
Physical simulation and verification. a. Mixture of 6 cell populations expression 3 orthogonal pairs of heterodimer surface binding pairs in K562 cell line. b. Based upon observation of microscopy data in ‘a’, a matrix of relative binding efficiency could be created. c. Physical conditions approximated by the physical model — cells are moving around a confined 3D space in Brownian motion. d. Simulation results for the number of cells present in ‘a’ with our model. e. Modeling and verification of a 2-tetrahedron shape. Upon applying the full modeling scheme to input shapes, the verification algorithm can compare simulated outcomes to the initial specification to determine frequency of success for this circuit. On the left is the desired shape and 3 simulated outcomes and their corresponding connectivity graph used to calculate verification score. f. Modeling and verification of the 3-tetrahedron shape. At the left is the specified shape and its corresponding connectivity graph. We show increasing scores (i.e. worse outcomes) of simulations from left to right. The scores are determined both for overall shape (left in brackets) and for individual tetrahedron formation (right in brackets).

At the beginning of the simulation, we create one cell, giving it the density of water and a radius of 6 microns and establish the ODE objects needed to represent the cell. This cell has a counter that counts how long it has been since it was last part of a division. If the cell is allowed to divide (i.e. it is not expressing the Cycle Arrest gene), then when the count reaches twenty hours the cell divides. New states are determined for the two daughter cells based on the state machine for the circuit.

As the cells divide and express proteins as per the state machine, the original cell becomes one of the daughter cells, while a second daughter cell is randomly placed a half cell radius away and apply a number of important physical considerations to the cells as they move around in the environment. First, we consider the forces applied to each individual cell in isolation. When unbound to another cell, a random force *F* = *F*_*x*_ + *F*_*y*_ + *F*_*z*_ is generated and then each cell receives a force *F* ^*′*^ = *r*_1_·*F*_*x*_ + *r*_2_·*F*_*y*_ + *r*_3_·*F*_*z*_ where *r*_1_, *r*_2_, and *r*_3_ are randomly picked for each cell from a uniform distribution on [0.5, 1.5). The correlated forces functionality was added to try to stop groups of cells from breaking apart too readily, and may in fact mirror reality more closely, given that there is no reason for connected cells to try to move in diametrically opposite directions. A drag force is also applied to the cells to try match the reality that the cells would not accelerate up to very high speeds in a liquid. The flow is assumed to be laminar (as opposed to turbulent) so the force applied is proportional to the cell’s velocity.

Second, we also consider distances between cells and change the forces applied if they are bound to other cells — when cells are too close, they repel each other, and when cells that are connected are too far apart, the connection either breaks or the cells are pulled together (to mirror the behavior the binder proteins). For this to occur, the ODE joints between the cells must be broken as the joints try to maintain their original lengths. Thus, the joints are broken, then reformed (unless the cells are too far apart) before the Brownian motion step. If a joint is no longer valid after a division — the two cells that were bound are no longer expressing the protein(s) that bound them — then the joint is removed. Or, if a new cell is near a cell that its parent cell was bound to, and that bond is still valid, then a joint is created between the new cell and the second cell. When two cells collide during the simulation the program checks to see if the cells can bind. If so, it creates a joint and changes their velocities according to a perfectly inelastic collision model.

Finally we use a mass spring optimization to correct cell distances for cells that are adjacent to each other in 4D space to add reality to the simulator. We use a compression of volume based model to repel cells if they are too close, and a spring model to draw cells together if they are bound and are too distant. For each cell, we first determine if cells are near each other. Cells are sorted into a 3D grid based on their coordinates. We traverse the grid, visiting cubes that have cells with them, calculating the distance between those cells and cells in at most thirteen neighboring cubes. We only need to consider the cubes in one half of the shell around each cube, as the other half will have already been visited by the algorithm. If the distance between any two cells is too close, the pair is added to a set. Finally, all cells that are bound to each other by surface binders are also added to the set.

Next we calculate the forces cells exert on each other due to either being too close (compression) or bound by surface binders but too far apart. If the cells are bound and are forced far apart, the joints between cells are broken. If the cells are too close, we calculate the fraction of their volumes that is shared, and adjust the position of cells to an optimal distance of 11 mircons. The volume of two overlapping sphere shaped cells *V*_*d*_, having an equal radius, at a distance *d* microns from their cell centers is define as:

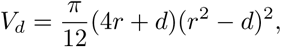

where *r* is the radius of each cell [44]. The optimal overlapping volume of two cells is defined as the shared volume of cells that have an optimal distance *d*_*o*_ of 11 microns between their cell centers:

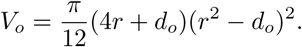

When cells become compressed as defined by having overlapping spherical boundaries, we can take the difference of the overlapping volume and the *optimal* overlapping volume to calculate a repulsive force that will push cells from their current distance to an optimal distance apart (11 microns). The force due to the compression *F*_*c*_ between two cells is calculated as:

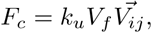

where *k*_*u*_ is the spring constant for pushing cells apart, *V*_*f*_ is the relative percent of volume overlapping of one cells (we call this the volume fraction), and 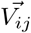 is a 3D unit vector pointing from the cell *i* to cell *j*. We can consider *k*_*u*_*V*_*f*_ as the magnitude of repulsion, and 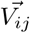 as the direction of repulsion. This force is equally applied to each cell that is too close to the current cell.

If the cells are too far apart (more than three cell radii), their joint is broken. Otherwise, a spring force is calculated — *F* = *k*_*p*_(*d* − 11) where *k*_*p*_ is a spring constant the pulls cells together and *d* is the distance between the cells. Once all the pairwise interactions have been considered, the forces due to the shared volumes are calculated.

As we show in (**Figure 7d**), once accounting for external events including cellular biophysics (**Figure 7c**) and orthogonal surface binding (both on- and off-target as per (**Figure 7b**), our simulations sufficiently approximate the forces of cell division, cell-cell adhesion, and Brownian Motion. This gives us confidence that as long as we model the internal genetic changes properly, the simulated spatio-temporal organization of the cell population should align closely with real cells in 4D (3D space + time) culture.

### 2.6 Verification of cellular growth outcomes

With the objective of enabling the programmatic assembly of cells into user-defined structures, a computational framework is needed to allow the user to specify a shape, design and build the required circuit, verify whether the specified shape is achieved for the candidate circuit, and determine the success rate of achieving the desired shape. Computer-aided verification allows us to test if a candidate circuit will result in the formation of the desired shape with a given probability and confidence level, eliminating the need to run many costly and time-consuming experiments.

In the verification step, we need to define a meaningful metric that captures different notions of the structure and design an algorithm to calculate this metric, with the goal to assign a quantitative value that can be used to compare the desired shape to the shape observed from a circuit. For this sub-problem, we use graphs to represent the desired and observed 3-dimensional structures. Each node in the undirected graph represents a cell, and the edge between two nodes demonstrates the cell-cell binding. We assume two input graphs are given in the verification step: the first graph *G*_1_ is produced from the CAD software in which the user identifies the desired shape and a graph is derived by applying meshing algorithms, and the second graph *G*_2_ is produced from the observed shape for a candidate circuit in our simulated model. The verification can be transformed into a graph matching problem that checks whether the two graphs *G*_1_ and *G*_2_ corresponding to the desired and observed shapes match. Therefore, classical graph matching algorithms [45] can be employed to solve this problem.

Graph similarity has been studied in various fields, from social networks [46] to biological networks [47], and many algorithms have been proposed to measure the similarity of graphs such as techniques using distance-based metrics and feature extraction [45]. We use the spectral eigenvalue similarity method [48] as the basis for our algorithm to determine the graph similarity, and propose an algorithm to calculate and score the similarity of the desired and observed graphs. Eigenvalues of a graph contain important information about its structure. For a given graph *G*, the degree matrix *D* is a diagonal matrix containing information about the degree of each node, i.e., the number of edges connected to the node. The adjacency matrix *A* is a square matrix with each element *a*_*i,j*_ indicating whether pairs of nodes *i* and *j* in the graph are adjacent *a*_*i,j*_ = 1 or not *a*_*i,j*_ = 0. The laplacian matrix *L* is defined as *L* = *D* − *A* and is a representation of the graph and can be used to define different properties of the graph structure (e.g. connectivity) and construct a low-dimensional embedding of the graph.

Let *L*1 and *L*2 be the laplacian matrices of graphs *G*1 and *G*2 with *n* nodes, respectively. The similarity score between the two graphs is defined based on the euclidean distance of the eigenvalues *λ* of the laplacian matrices as:

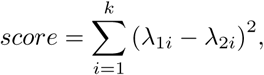

where *score* is a real non-negative number, *λ*_*ji*_ is the *i*^*th*^ largest eigenvalue of the graph *j* and *k* is chosen such that the top *k* eigenvalues contain 90% of the graph spectrum energy:

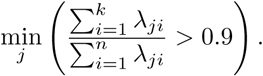

The eigenvalues of the laplacian matrix of a graph are invariant with respect to node permutations. Therefore, if two graphs *G*1 and *G*2 are isomorphic, i.e., the desired and observed shapes are exactly the same and possibly rotated, their laplacian matrices will have the same eigenvalues, and the *score* will be 0. A *score* closer to 0 indicates that graphs are more similar, while higher values show dissimilarity of the graphs.

In the considered cases the desired structure consists of substructures (blocks of tetrahedron) of cells with different expressions (colors). Therefore, we employ a 2-stage algorithm where the similarity score is determined by the similarity of the overall observed structure with the desired one, augmented with the similarity of each substructure compared to a single block (tetrahedron). We use an upper-bound threshold ∊ for which we can automatically discard simulations with a similarity *score* greater than ∊, i.e., we reject the circuits which yield to dissimilar shapes. Another challenge here is the fact that the observed graphs (shapes) for a given circuit can possibly have different number of nodes (cells) and edges (binding connections). To tackle this issue, we measure the similarity by comparing the desired graph with the largest connected subgraph in the observed graph. We also provide the expected probability of achieving the desired shape and determine the statistical significance of the candidate circuit.

For the two example shapes in (**Figure 7e,f**), we simulated 96 outcomes from one starting cell, resulting in a variety of scores, which is expected for this stochastic simulation process. Fortunately we saw many shapes that both appeared to be the 2-tet and 3-tet shape we wanted and got strong scores from the verification algorithm. This algorithm helps us sort through massive amounts of image to determine if we are successfully forming shapes. Since their are many different types of errors that can occur resulting in different scores that are all in relative units, a scientist must draw a line for where success is measured to determine the ultimate chance of success of a given shape.

In general, the proposed graph matching algorithm provides reasonable results for the case studies we consider in this paper. More advanced techniques can be employed to verify the similarity of more complex shapes. One approach is to define other shape metrics such as sphericity, as well as the graph similarity score, and apply machine learning methods to cluster (unsupervised) or classify (supervised) large sets of simulated shapes.

## 3. Discussion

This work describes a computer-aided design (CAD) software, *CellArchitect*, that can be used to create genetic circuits that can be integrated into one cell and to grow into arbitrary multi-cellular 3D shapes. It represents the first described CAD workflow for solving the developmental biology problem of shape formation entirely based on genetic circuits. While we show some simple examples end-to-end to demonstrate that each sub-problem solution is reasonable, the software is built to accommodate shapes of arbitrary size. Each individual sub-problem solution could likely be improved — however, we are the first to break the multi-cellular shape formation into solvable sub-problems with clear boundaries: 1. using a geometry to define substructures; 2. creating developmental trees to represent which cellular progenitors should be responsible for certain tasks; 3. developing genetic circuits to count cell cycles and express genes at discrete developmental time points; 4. using a finite-state-machine model to simulate genetic recombination circuits over time from a single-cell origin; 5. simulating cell growth in the context of internal (genetic circuit) behavior and external (environmental) variables in 4D; and 6. using quantitative verification methods to compare the desired shape to the shapes observed both *in silico* and *in vitro*. This method or others like it will likely be needed to execute complicated synthetic developmental behavior — human cells simply take too long to grow for trial-and-error approaches to be debugged on any reasonable time scale.

While we defined sub-problems that must be solved for shape formation and provided solutions for each of these areas, there are some limitations of these solutions that must be built on in future work to allow this software to scale properly. First, the state-machine solution does not scale well with increasingly complicated genetic circuits, so we choose to ignore rare events. Even in this case, the computational cost of making these determinations gets very high — this will need to be addressed for larger cells masses. Similar problems exist for the simulation framework — the cell-cell interactions complexity also scales exponentially. These bio-physical and genetic assumptions will need to be revisited and improved upon in future versions of this software.

Some sub-problem solutions are also potentially useful for other applications or future projects in different offshoot projects. Specifically, the developmental tree is a potentially interesting idea in problems not specifically related to shape formation. Many different types of events happen during development, such as the expression of certain signaling factors. Mapping back cell division events and identifying asymmetrical events and when cells stop proliferating. This model might be useful for those types of problems as well. Our developments in automated genetic circuit design for developmental biology might also be useful for other applications. As with the developmental tree, it would be potentially useful for problems outside of shape formation, such as the inclusion of genes to change cell type. It is already an open challenge to get cell masses of heterogeneous cell type composition, so using our counter constructs to signal differentiation and asymmetrical division during specific time points in development could be useful outside of shape alone. Furthermore, our developments in adding the first known framework for simulating the impacts of a genetic circuit over time based upon component characterization data could be applied to recombinase-based circuits outside the context of human or developmental biology. Combined with the automated genetic circuit design, these two modules could be applied in many different types of genetic circuits in different types of organisms to simulate outcomes of system-level behavior with synthetic biology circuits.

While we have attempted to create a framework for solving the challenging problem of shape formation, the larger idea was to develop a new framework for thinking about ‘synthetic developmental biology’. While there are certainly other exciting approaches to scaling up tissue engineering and developmental biology applications, we have added a new framework for genetics and synthetic biology to synergize with other complementary technologies. For example, 3D bioprinting might be a good tool to form larger structures with populations of cells, but it still might be important to genetically form certain small-scale structures over time in a tissue. While we generally overlook complex media condition manipulations in our current work, all cells must grow in media, and so future extensions to include considerations for this could be highly beneficial. This work also complements well with existing work to differentiate cells with transcription factors — the combination of the ability to change cell type and multi-cellular shape autonomously in the same genetic system could lead to the creation of novel tissue-like structures that might be hard to create with tools like 3D-bioprinting.

## 4 Methods

All algorithms and software are implemented in Python 3 using numpy, panda3d, matplotlib, dnaplotlib, and psutil. The cell simulator uses Open Dynamics Engine (ODE) as a physics engine, as implemented by the panda3D package (note that the underlying ODE framework is implemented in C++). Cell simulation visualization is performed in ParaView. Large computing jobs were carried out on a Linux-based computing cluster.

## 5 Acknowledgments

EA, TD, MM, GC, and TW developed meshing and genetic circuit software components. EA, NM, TD, DB, IH, ACP, and CB developed the simulation and verification software. GC and TW provided data for physical simulation parameters. EA, GC, TW, and GC devised the preliminary vision of the project. EA, NM, TD, DB, CB, and GC wrote the paper. The authors would like to thank Justin Gallivan, Jesse Dill, Blake Bextine, Nathaniel Borders, Clair Travis, and Helene Kuchwara for helpful conversations. This work was supported by the DARPA ELM Program under contract W911NF-17-2-0079.

